# Weakly supervised learning uncovers phenotypic signatures in single-cell data

**DOI:** 10.1101/2024.07.29.605625

**Authors:** Anastasia Litinetskaya, Soroor Hediyeh-zadeh, Amir Ali Moinfar, Mohammad Lotfollahi, Fabian J. Theis

## Abstract

To deliver clinically relevant insights from large patient cohorts profiled with single-cell technologies, a key challenge is to relate sample-level and single-cell measurements. We present MultiMIL, a deep learning framework that applies attention-based multiple-instance learning for phenotype prediction and cell state identification. We applied MultiMIL to peripheral blood mononuclear cells from COVID-19 patients, the Human Lung Cell Atlas, and a spatial proteomics breast cancer dataset, demonstrating how our model can be utilized to find phenotype-associated cell states, learn phenotype-informed sample representations, and expand disease signatures.

## 1 Background

The advancements in single-cell technologies have enabled a consistent increase in the number of cells captured for analysis, as well as the number of samples, i.e., individual donors, tissue biopsies, or other patient-level observations, resulting in larger cohort sizes in atlases [1, 2]. To understand the underlying cellular processes and mechanisms that drive disease phenotypes, a significant challenge remains in linking cell-level signals to patient-level phenotypes in an interpretable manner.

Multiple strategies have been developed for this task, including pseudobulking, which aggregates cell-level data to mimic bulk profiles and facilitates sample-level analysis; methods that identify cell types exhibiting differential transcriptomic signals (e.g., via compositional shifts or marker expression); and direct phenotype prediction models operating at either the cellular or patient level [3–14]. However, these approaches often suffer from limitations. Pseudobulking sacrifices single-cell resolution; many methods are restricted to scRNA-seq and do not support other modalities. Additionally, some methods struggle to disentangle genuine biological variation from technical or batch effects. Even when predictions are made at the sample level, it is not straightforward to link these predictions to the cellular processes driving the disease phenotype.

In machine learning terms, predicting sample-level phenotypes from cell-level data is a weakly supervised learning problem—only sample labels are available, while labels for individual cells are missing. Multiple-instance learning (MIL) offers a suitable framework: each sample (bag) contains multiple cells (instances), and the goal is to learn bag-level labels while identifying which instances are most informative [15]. Attention-based MIL, in particular, enables interpretable aggregation by assigning weights to each cell’s contribution to the sample-level decision. MIL has been successfully applied in various biomedical domains, including blood disorder classification from microscopy, learning genetic variant associations with transcriptional states, and T-cell receptor sequence-based cancer detection and phenotype prediction from scRNA-seq data through hierarchical MIL frameworks that incorporate cell-type information [16–19].

When applying MIL to single-cell data, several domain-specific challenges must be considered. Biological signal is often confounded by technical batch effects and measurement noise, complicating the task of distinguishing true disease-associated variation from artifacts. From a machine learning perspective, it is challenging to balance predictive performance with interpretability, as highly predictive models often do not perform robustly enough to rely on for hypothesis generation. For the approach to be helpful in diagnostics or therapeutics, the identification of disease-associated cellular states must be reproducible, indicating that they drive disease phenotypes rather than reflect noise or artifacts.

To address these limitations, we introduce MultiMIL, a deep MIL-based model for phenotypic prediction and cell prioritization. Since not all cells from diseased samples exhibit disease-induced changes, the MIL approach enables the identification of cells most affected by the disease. Identification is achieved through attention scores, where highly attended cells are most relevant and the least attended cells are the least relevant to the observed phenotype of the sample. This reflects the two main mechanisms — transcriptional shifts or changes in cell type composition — through which the observed phenotypes can be explained. MultiMIL is agnostic to the input modality and can be trained on raw features or latent representations from atlases or foundation models.

We demonstrate the power of MultiMIL’s phenotypic prediction for unseen patients and prioritization of disease-specific cell states by analyzing several high-relevance clinical datasets: a human peripheral blood mononuclear cells from COVID-19 patients profiled with single-cell RNA-seq [20], the Human Lung Cell Atlas [21] and a breast cancer dataset, profiled with imaging mass cytometry [22]. We further demonstrate how the disease states identified with MultiMIL help discover novel genes associated with the disease and finally, how the sample representations learned with our model are reflective of the phenotypes. MultiMIL’s implementation and reproducibility scripts are available at http://github.com/theislab/multimil. We also publish a pipeline for comparing different methods for sample-level prediction from single-cell data at https://github.com/theislab/sample-prediction-pipeline, which enables fast, multi-scale analysis of custom datasets.

## 2 Results

### Prioritizing phenotype-specific cells and predicting phenotypes with MultiMIL

MultiMIL is a deep-learning-based model that allows the prediction of sample-level phenotypes from single-cell measurements and the identification of cell states associated with these phenotypes. MultiMIL’s model consists of a pooling layer that learns sample-level representations from single-cell representations and a classification network that learns to predict sample-level phenotypes (**Fig. 1b**). We draw inspiration from the multiple-instance learning (MIL) approach [15, 16], where we model samples as bags and cells as instances belonging to a bag. The classification labels are only known on the bag level but not on the instance level, and we are interested in identifying instances associated with the bag label (**Suppl. Fig. 1**).

**Figure 1.**
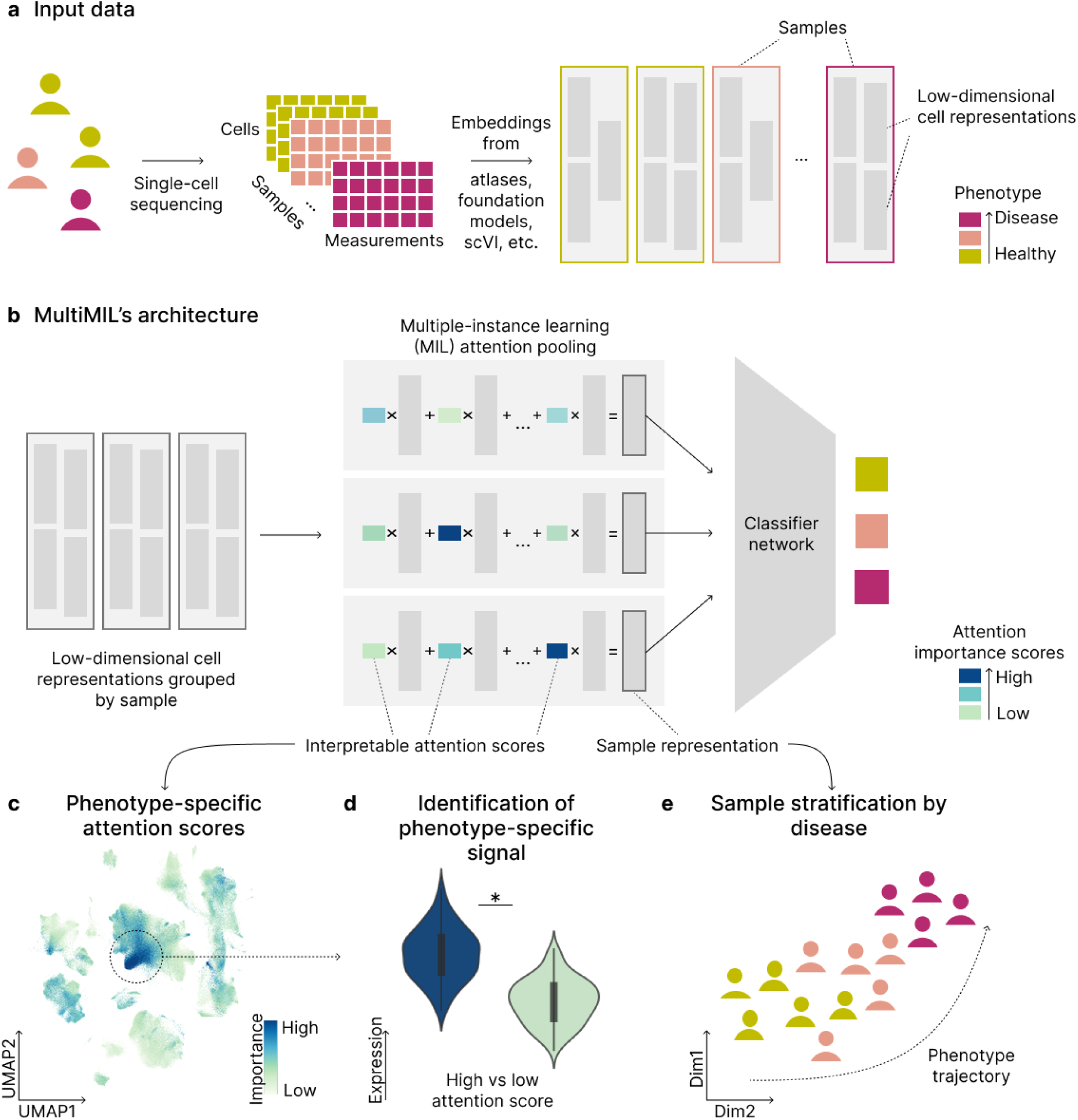
MultiMIL predicts sample-level phenotypes through interpretable multi-scale learning. **(a)** MultiMIL takes low-dimensional cell representations grouped by sample as input. **(b)** MultiMIL’s architecture consists of a MIL attention pooling layer, which takes as input cell representations and outputs sample representations, and a sample classification network that predicts phenotypes from sample representations. **(c)** MultiMIL learns an attention score per cell to identify cell states associated with phenotype and **(d)** allows for novel differential signal identification. **(e)** The learned sample representations reflect the phenotype trajectory.

The input to the model is usually batch-corrected cell embeddings, for instance, embeddings from atlases or foundation models (**Fig. 1a**). It is also possible to train MultiMIL directly on the raw features. The MIL attention pooling layer aggregates the cell-level embeddings into a bag embedding employing attention pooling. During training, the model learns attention weight *α_i_* for each cell *i* in a bag and then aggregates cell embeddings *z_i_* into a bag representation *z*_bag_ as a weighted sum z_bag_ = Σ*_i_ α_i_z_i_*. The pooled representation *z*_bag_ is then fed into a feed-forward classification network that predicts the phenotypes.

MultiMIL provides several ways to interpret the learned attention weights (**Fig. 1c**). Firstly, the higher the weight of a particular cell is, the more important the cell is for the prediction. By learning a score for each cell, we identify and analyze cell states associated with a particular phenotype by selecting cells with high attention scores. This allows for the identification of differential signals between groups at higher resolutions than cell types, as we are able to subset the data to only disease-affected cells (**Fig. 1d**). Attention scores also allow us to learn phenotype-informed sam-ple representations. We can obtain them by averaging the cells’ representations with the highest attention scores within a sample (**Fig. 1e**).

MultiMIL is fast to train due to the mini-batch training and the deep-learning nature of the model: the model training for the dataset of approximately 250,000 cells takes less than 6 minutes (Table 3). Computational efficiency, combined with interpretability through attention scores, makes Mul-tiMIL a scalable and practical tool for identifying disease-relevant cellular signals in large single-cell datasets.

### Robust phenotype prediction from diverse single-cell embeddings with MultiMIL

In this section, we will investigate how the quality of the embedding affects the prediction performance and the consistency of the learned attention scores. We will also benchmark the model’s performance depending on various training and model parameters such as batch size, learning rate or pooling function (the range of tested parameters is shown in Table 1 in Methods). We used a peripheral blood mononuclear cell (PBMC) dataset [20] throughout these experiments. This large-scale dataset consists of 130 healthy and diseased samples and provides annotation on the progression of COVID-19 stages.

**Table 1.** Hyperparameter search for MultiMIL’s prediction.

| Hyperparameter | Description | Default | Range |
| --- | --- | --- | --- |
| Learning rate | learning rate parameter | 1e-4 for single-cell, 1e-3 for IMC | $\{1e-5, 1e-4, 1e-3, 1e-2\}$ |
| Batch size | number of cells in a forward pass | 256 | $\{256, 512, 1024\}$ |
| Pooling function | how the sample representations are calculated | gated attention | $\{\text{gated attention, attention, mean, max}\}$ |
| Epochs | number of epochs the model was trained for | 50 | $\{50, 100, 150, 200\}$ |

First, we investigated how the quality of the embedding and different model parameters affect the performance of MultiMIL. To this end, we obtained low-dimensional representations of the RNA-seq single-cell measurements using scGPT and GeneFormer foundation models. We also trained an scVI model and calculated the PC embeddings before batch correction as two additional baseline embeddings. We subset the data to healthy, mild and severe COVID-19 samples, split the data into five cross-validation splits and train MultiMIL to predict these 3 classes either as a classification task or an ordinal regression task. We observed that the classification accuracy was correlated with the quality of the embedding, which we assessed with scIB integration metrics [23]. The accuracy was the highest for scVI and scGPT embeddings (83% and 84%, respectively, averaged for classification and regression) and lower for PC and GeneFormer embeddings (80% and 79%, respectively) (**Fig. 2a**, **Suppl. Fig. 2b, c**). The regression model consistently outperformed the classification model on the 3-class prediction problem (**Fig. 2a**), and we observed that the variance of the accuracy was also consistently lower for the regression model (**Suppl. Fig. 2a**). However, the regression model performed worse than the classification model on the 6-class task. On the task of binary prediction, classification and regression models performed similarly, indicating that for easier tasks, both models can perform equally well (**Suppl. Fig. 3d**). We additionally compared the default classification and regression models to the alternative architecture where we concatenated one-hot encoded embeddings for sample covariates to the sample embedding before passing it through the classification network (see Methods). Incorporating the covariates decreased the performance accuracy (**Suppl. Fig. 3e**), so we used the model architectures without including the additional sample-level information.

**Figure 2.**
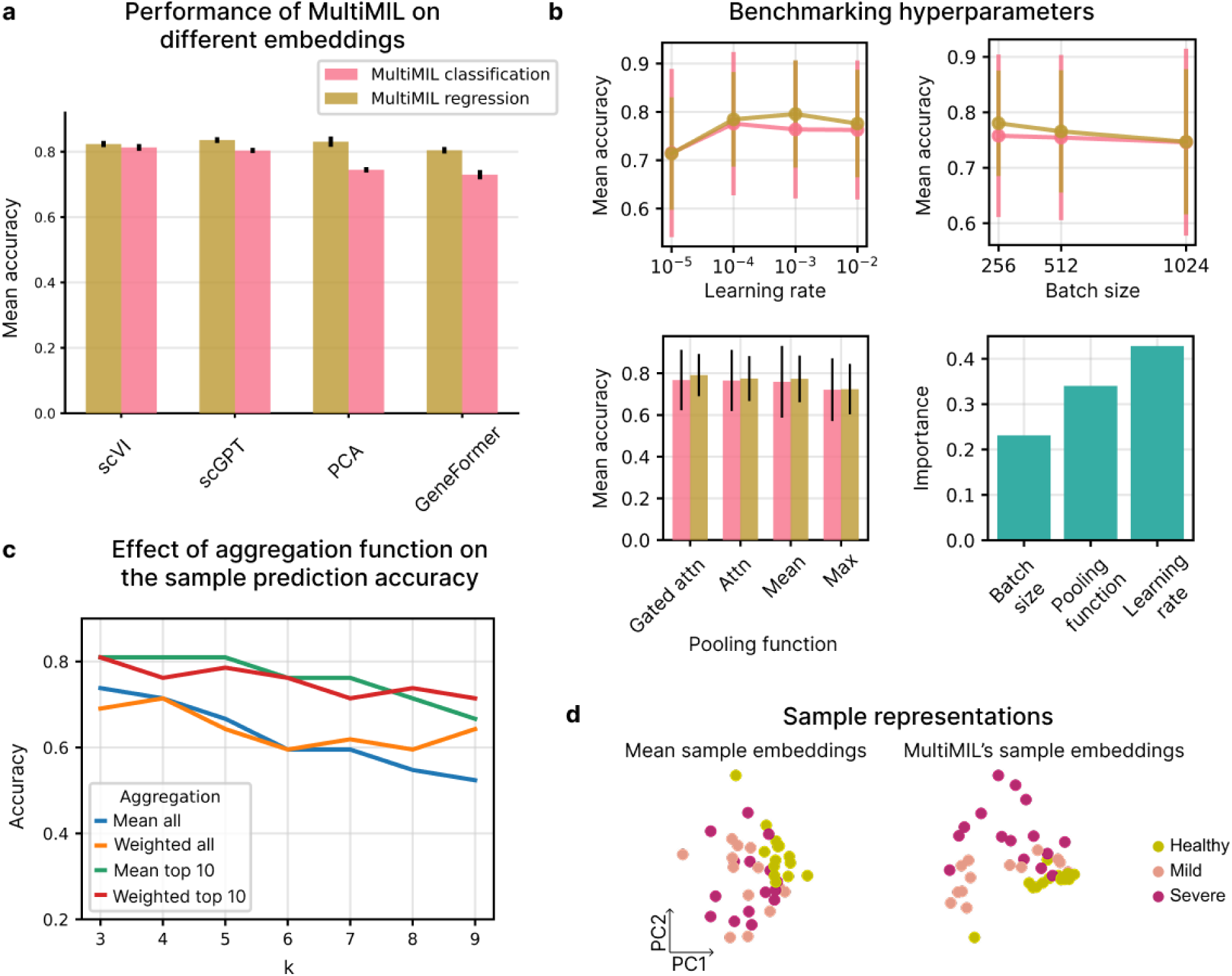
Benchmarking MultiMIL. **(a)** Best accuracy achieved by MultiMIL’s classification and regression models on different embeddings. **(b)** Effect of the learning rate, batch size and the pooling function on the accuracy and the feature importance score of these parameters. **(c)** Sample-level kNN prediction accuracy of different sample-representation aggregation methods. **(d)** PC plot of mean sample embeddings and MultiMIL’s sample embeddings colored by disease stage.

Next, we investigated the effect of the learning rate, batch size and the pooling function on the performance, focusing on the better-performing embeddings, i.e., scVI and scGPT. We also trained a random forest regressor to find which hyperparameters influence the performance most. Gated attention pooling slightly overperformed the mean pooling and the attention pooling, while max pooling consistently underperformed (**Fig. 2b**). Since we want to benefit from the interpretability of the attention scores, we use gated attention as the default for the rest of the analyses. We observed that the batch size did not have a significant effect on the performance, although the overall stability of the predictions was slightly higher for smaller batch sizes (**Suppl. Fig. 2d**). Learning rate was the most important hyperparameter for the accuracy (**Fig. 2b**), so we recommend that users start with a learning rate optimization if the default parameters yield unsatisfactory performance. We also observed that training the model for 50 epochs yielded the best performance, compared to longer training regimes (**Suppl. Fig. 2e**).

We also investigated how consistent the cell attention scores are across cross-validation splits, comparing the results from best-performing models trained on scVI and scGPT embeddings. We observed that the model trained on the scVI embeddings consistently learned high attention for the same cell types (**Suppl. Fig. 3a,b**), so we hypothesize that the dimensionality of the latent space could play a role, where the input of lower dimensionality yields more consistent cell attention scores (30 dimensions for scVI vs. 512 for scGPT).

Finally, we show that by aggregating cells with the highest attention scores, we obtain sample representations most indicative of the disease stages, compared to averaging all cell embeddings or taking a weighted (by attention score) average of the cell embeddings (**Fig. 2c,d**). Averaging cell embedding of the top 10% cells with the highest attention score yielded the best performance, compared to other thresholds (**Suppl. Fig. 3c**).

We showed that MultiMIL’s performance benefits from good-quality cell embeddings. The model can robustly predict sample-level covariates across different hyperparameters and aggregation strategies.

### MultiMIL accurately predicts COVID-19 stages and identifies disease-associated cell states

Next, we discuss the results on the PBMC dataset [20], comparing MultiMIL’s predictive performance to baselines and linking the high-attention cell states to known COVID-19 pathology. We used the scVI embedding subset to healthy, mild and severe COVID-19 samples and selected the best-performing MultiMIL’s model on this 3-class prediction task. We assessed the predictive performance of MultiMIL compared to several baseline models and interpreted the cell attention scores (**Fig. 3a**).

**Figure 3.**
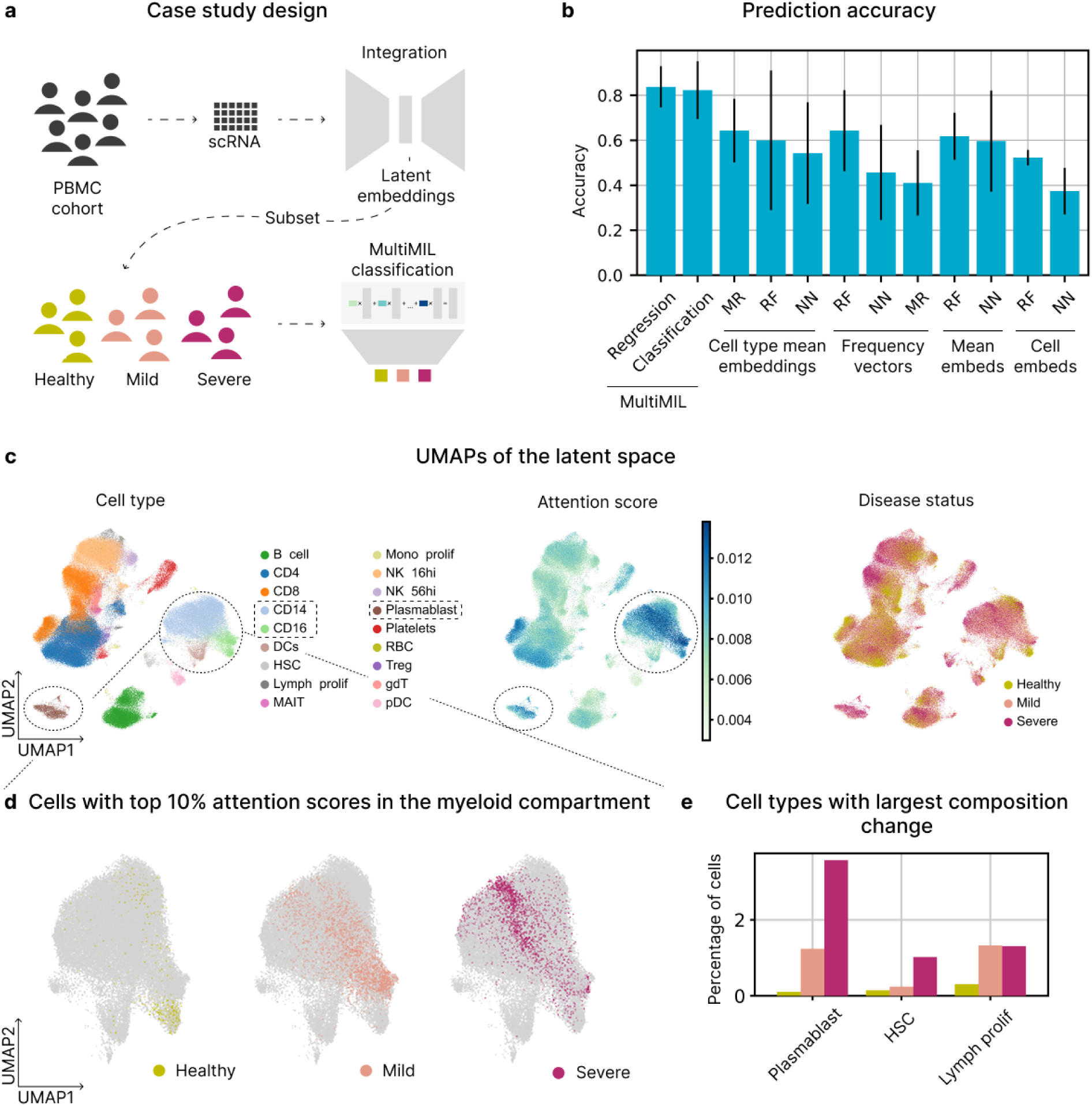
MultiMIL accurately predicts disease stages on a PBMC dataset. **(a)** Case study design. PBMCs were sequenced with CITE-seq (paired RNA and ADT), integrated to obtain a shared latent representation, subset to healthy, mild and severe COVID-19 samples, and used as input to train MultiMIL’s classification network. **(b)** A bar plot showing average accuracies and standard deviations (i.e., the length of an error bar equals two standard deviations) of the five cross-validation runs on the disease-prediction task. MultiMIL was trained in the classification and regression settings. Cell-type mean embeddings and frequency vectors were input to the random forest (RF), feed-forward neural network (NN) and multiclass logistic regression (MR) models. Mean embeddings and cell embeddings were input to the RF and NN models. **(c)** UMAPs of the integrated latent space colored by cell type (left), cell attention scores (middle) and condition (right). The myeloid compartment (i.e., CD14, CD16 monocytes and dendritic cells) and plasmablasts have high attention scores. **(d)** UMAPs of the myeloid compartment showing the healthy, mild and severe COVID-19 cells with the top 10% of attention scores for each condition. **(e)** A bar plot showing the top three cell types with the biggest compositional change from healthy to severe COVID-19, including plasmablasts.

In this work, we focus on weakly-supervised prediction where the sample-level labels are known, so we compared our model to supervised baseline models, including a random forest, feed-forward neural network and multiclass regression. Approaches utilizing single-cell data for phenotypic prediction often rely on (pseudo-)bulk data [7, 24], so we included a range of pseudo-bulk baselines in our comparison. Since MIL models generally fall between models that make predictions on the instance (i.e., single-cell) level and models that make predictions on the sample (i.e., bulk) level, we also include cell-level baselines (**Fig. 3b**, Methods). The mean embedding of a sample is the mean of cell embeddings belonging to this sample, and cell-type mean embeddings are calculated as the mean of cell embeddings per cell type and concatenated per sample. Frequency vectors are calculated as relative frequencies of cell types present in each sample. For cell embeddings, the input to the models was the cell embeddings from the scVI space, and the prediction was made for each cell. We note that cell-type mean embeddings and frequency vectors are supervised since the cell type labels are required, while MultiMIL, mean embeddings and cell embeddings are not. MultiMIL outperformed all the baselines in a 5-fold cross-validation experiment, achieving an accuracy of 84% (**Fig. 3b**).

When analyzing diseased samples, we are interested in identifying cell states affected by the disease. By utilizing the attention pooling layer, our model learns a weight for each cell, where higher weights directly correspond to cell states associated with the condition. We only take into account cells with the 10% highest scores per condition, as these cells are most strongly associated with the disease. We observe in **Fig. 3c** that cell types with the highest attention scores are CD14 and CD16 monocytes. Hence, we first examine the myeloid compartment (**Fig. 3d**) and notice a trajectory of highlighted CD14 monocytes from healthy and mild to severe, indicating a shift in expression levels between different stages. Similarly, we find distinct populations of highlighted healthy and mild CD16 monocytes, confirming that the signal learned with MultiMIL aligns with previous studies reporting strong changes in monocytes with the progression of COVID-19 [25, 26].

We also noticed that the plasmablast cluster had a high attention score, so we hypothesized that it could be related to compositional differences. Hence, we next investigated which cell types had the biggest compositional changes between conditions and found that plasmablasts was the cell type most affected between healthy and severe conditions (**Fig. 3e**). MultiMIL identified these compositional changes as indicative of disease progression, also reported in [20]. We additionally ran Milo [13] on the same embeddings and found that cell populations identified by MultiMIL, e.g., CD16 monocytes and platelets, were among the cell types with the highest log-fold-change in composition identified by Milo (**Suppl. Fig. 4**). We note that Milo allows comparisons between two conditions, while MultiMIL identifies condition-specific cell states for multiple classes simultaneously.

These findings reinforce the role of classical and non-classical monocytes in COVID-19 severity progression and demonstrate that MultiMIL can rediscover key immune modulators of disease progression. The ability to distinguish myeloid shifts between mild and severe cases without using cell type labels highlights MultiMIL’s utility for studying immune responses.

### MultiMIL identifies a subpopulation of IPF-associated macrophages in human lung

Single-cell atlases provide integrated and cell-type-harmonized representations of different systems or organs of interest. These atlases can comprise hundreds of donors, which in turn is crucial to understanding the disease variability and potential therapeutic targets [27]. We demonstrate how MultiMIL can be utilized with existing single-cell atlases. The Human lung cell atlas (HLCA) [21] consists of healthy and diseased donors integrated into a common latent space. We investigated idiopathic pulmonary fibrosis (IPF) and compared diseased and healthy samples. We selected healthy and IPF donors from the atlas and trained MultiMIL in a 5-fold cross-validation setting (**Fig. 4a**). On this classification task, MultiMIL outperformed all baselines, although several other models also achieved strong performance, with accuracies exceeding 90% (**Suppl. Fig. 5d**). Notably, if the primary goal is binary classification rather than interpretability, simple mean-embedding approaches can still offer competitive results.

**Figure 4.**
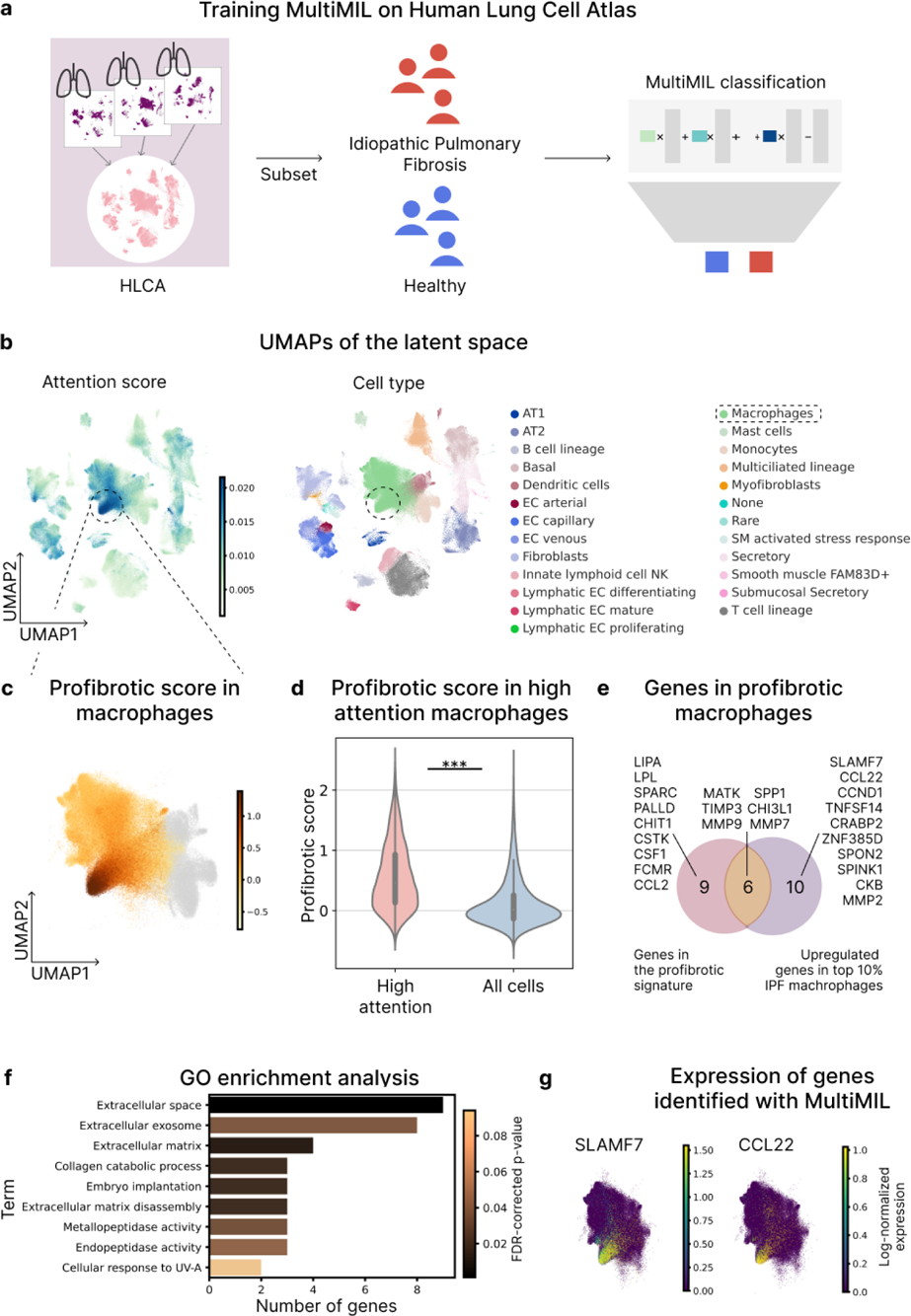
MultiMIL identifies IPF-associated cell states in human lung macrophages. **(a)** Pre-trained embeddings from the HLCA were subset to healthy and IPF samples and used to train MultiMIL’s classification network on the binary classification task. **(b)** UMAPs of the original latent space from the HLCA colored by cell type (top), condition (bottom left), the profibrotic score calculated in macrophages (bottom middle) and cell attention score (bottom right). A subpopulation of macrophages has a high attention score, so we investigate these cells further. **(c)** A UMAP plot of macrophages colored by the profibrotic score. **(d)** Violin plots showing the profibrotic score in high-attention macrophages and all macrophages from IPF patients (p-value<0.001, two-sided t-test). **(e)** A Venn diagram with the genes in the profibrotic signature, the number of genes that are upregulated in the high-attention macrophages compared with all macrophages from IPF patients and the number of genes in the intersection of the two sets. **(f)** GO enrichment analysis of the upregulated genes in the high-attention macrophages. **(g)** UMAPs of the macrophages colored by the expression of *SLAMF7 and CCL22*.

We examine the learned cell attention scores to analyze which cell states the model learns to associate with the disease. We notice that a subset of macrophages has the highest scores (**Fig. 4b**), so we first show that MultiMIL identifies a subpopulation of *SPP* 1^hi^ IPF-specific macrophages [5, 28] (**Suppl. Fig. 5a**). We hypothesize that this subpopulation corresponds to profibrotic macrophage populations reported in previous studies [29, 30]. To confirm, we calculate the profibrotic score based on the profibrotic signature introduced in [29] (**Fig. 4c**). We select macrophages from IPF patients and show that the cells with the highest attention score (top 10%) have a significantly higher profibrotic score than all IPF macrophages (**Fig. 4d**). MultiMIL also identifies a KRT17^+^ subpopulation of basal cells (**Suppl. Fig. 5b**) that previously have been reported to be associated with IPF [5, 31].

Cells with high attention can also be used for novel gene signature discovery or to expand the existing signatures. We demonstrate how to identify the gene signature of the IPF-associated macrophage subpopulation using only the attention scores and not relying on previous knowledge. We ran edgeR [12] to find differentially expressed genes between IPF macrophages with the top 10% highest attention scores and all IPF macrophages and identified 16 significantly upregulated genes. Comparing these 16 genes with the genes from the profibrotic signature, we find the overlap of 9 (out of 15) genes (**Fig. 4e**).

The genes identified solely from MultiMIL’s high attention group include *SLAMF7*, which has been previously reported to regulate the immune response in lung macrophages during polymicrobial sepsis and COVID-19 [32, 33]. Elevated levels of *CCL22* have also been found in patients with IPF [34, 35]. *TNFSF14* promotes fibrosis in the cardiac muscle and atria [36], lung [37] and kidney [38]. Interestingly, *TNFSF14* has been reported to regulate fibrosis in both structural and immune cells [37] (**Fig. 4g**, **Suppl. Fig. 5c**).

IPF is driven by excessive extracellular matrix (ECM) deposition and a disrupted balance between ECM production and degradation, processes in which the matrix metalloproteinase (MMP) and tissue inhibitor of metalloproteinase (TIMP) systems, particularly in macrophages, play a central role [39]. Among the top markers identified by our differential expression analysis were well-established profibrotic genes, including *TIMP3, MMP7*, and *MMP9*, all previously linked to IPF pathology. In addition, we uncovered a set of genes—*CCND1, CRABP2, SPON2, SPINK1, CKB*, and *MMP2* —with known roles in ECM remodeling and fibrosis-related pathways [40–45]. To further support the functional relevance of these findings, Gene Ontology enrichment analysis [46, 47] of the 16 upregulated genes in the high-attention group revealed a strong enrichment for ECM-associated processes (**Fig. 4f** ).

By identifying the subset of profibrotic macrophages previously implicated in ECM remodeling and fibrotic progression, MultiMIL independently validates known pathogenic populations and suggests additional genes that may contribute to fibrotic signaling, thus expanding current disease gene signatures.

### MultiMIL predicts grades in breast cancer from IMC data

MultiMIL can also be utilized directly with the raw measurements of an imaging mass cytometry (IMC) experiment and map learned attention scores to the spatial domain. We use a dataset of pathology images of breast cancer tumors [22] of different subtypes (hormone receptor (HR) +/-, HER2 +/-) and grades (1-3). The panel of 35 antibodies included breast cancer-specific biomarkers like estrogen receptor (ER), progesterone receptor (PR), HER2, the proliferation marker Ki-67, as well as lineage markers. Some of the present biomarkers were shown to be associated with higher grades, e.g., Ki67 or Vimentin [48, 49], so we focused our experiments on the grade prediction.

We trained MultiMIL to predict tumor grades (1-3) and achieved an accuracy of 0.87, substantially outperforming all baseline models, with the next-best accuracy of 0.74 from a random forest trained on cell-level input (**Fig. 5a, Suppl. Fig. 6a**). To interpret these predictions, we examined the cells with the highest attention scores and their spatial distribution. Most of these high-attention cells were tumor cells (**Fig. 5b,c**), indicating that the model relies heavily on tumor cells to distinguish between grades. Interestingly, the top-attention cells were consistently located in the tumor core rather than at the periphery. A co-occurrence analysis [50] confirmed that the model preferentially attends to tumor cells surrounded by other tumor cells (**Fig.5d**), a pattern observed across all three grades (**Suppl. Fig. 6b**).

**Figure 5.**
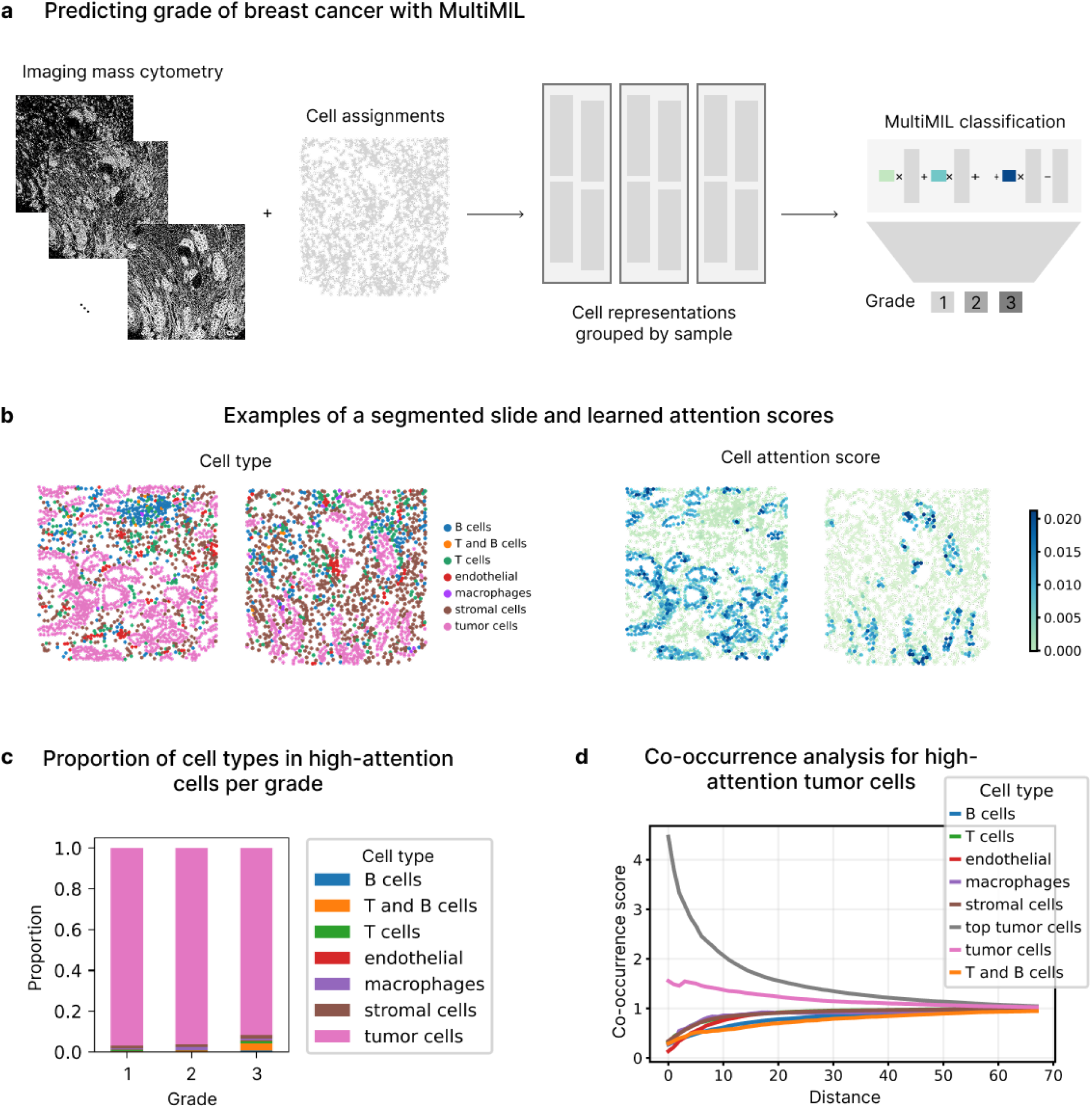
MultiMIL identifies spatially co-localized grade-specific tumor cells in IMC breast cancer data. **(a)** MultiMIL was trained to predict grades of breast cancer from the channels of IMC data, segmented to the single-cell resolution. **(b)** Two example slides colored by cell type and attention score. **(c)** Bar plot showing the proportion of cell type among high-attention cells per grade. **(d)** Co-occurrence of high-attention tumor cells with other cell types averaged across grades, depending on the distance on the slide.

Next, we examined how high-attention tumor cells differ from lower-attention ones. We performed differential expression testing within each grade and identified several biomarkers previously reported as dysregulated in breast cancer. For instance, GATA3 was upregulated in high-attention tumor cells in grades 1 and 2 [51], while Vimentin and CD44 were enriched in grade 3 [52, 53]. To further test whether focusing on the most informative tumor cells enhances detection of grade-specific signals, we restricted the analysis to top-attention tumor cells and observed increased sensitivity: for grades 2 and 3, this approach revealed more differential biomarkers (3 and 8 vs. 4 and 11, respectively; Suppl. Table 1). Notably, p53 showed stronger differential expression between grades when restricting to top-attention tumor cells (1.5 log-fold change) compared to all tumor cells (1.26 log-fold change). Interestingly, studies have reported conflicting associations between p53 expression and tumor grade: while one study found no association between p53 and tumor grade in patients with visceral metastatic breast cancer [54], another showed that p53+ status correlated strongly with grade 3 tumors and basal subtype in African-American women [55]. This discrepancy may reflect underlying heterogeneity, where only a subset of tumor cells contributes to the observed associations. Our findings support this view and suggest that these informative subpopulations highlighted by MultiMIL may correspond to spatially co-localized tumor cells in the tumor core.

## 3 Discussion

MultiMIL provides a deep-learning framework for predicting sample-level phenotypes from single-cell data while simultaneously identifying disease-associated cellular states. By combining attention pooling with multiple-instance learning, our approach directly links cell-level features to sample-level outcomes, offering a scalable pipeline for atlas-scale analyses. Unlike classical aggregation strategies, MultiMIL highlights the cellular populations most predictive of disease and pinpoints the transcriptomic and compositional changes that drive these differences. Additionally, MultiMIL can be utilized to predict phenotypes for unseen samples and learn sample representations.

Multiple-instance learning has proven effective for linking information across scales. In pathology, it has been widely used to connect cellular features with whole-slide level predictions [56, 57]. Early studies have begun adapting MIL to single-cell data, either in imaging-based or genomics contexts [16, 17]. Here, we extend this idea by framing single cells as instances and biological samples as bags, demonstrating how MIL can provide a framework for analyzing single-cell genomics. Future work should systematically benchmark MIL-based approaches to identify optimal strategies for different single-cell applications.

As with other deep-learning methods, MultiMIL is prone to variability due to stochastic training and hyperparameter sensitivity. We found that attention-based cell state prioritization can fluctuate across runs. To mitigate this, we recommend training across multiple splits and interpreting aggregated attention scores. Hence, attention weights should be viewed as hypothesis-generating signals rather than definitive markers of causal biology.

The growing availability of large-scale resources such as the Human Cell Atlas [1, 58] makes MIL-based approaches especially timely. As more large-scale atlases are released, MultiMIL can be readily applied to these datasets to identify cell states potentially relevant to various diseases. This will be particularly impactful in complex diseases such as Alzheimer’s, where large cohort datasets are already available [59, 60]. MultiMIL will facilitate the discovery of novel disease-associated cell states and mechanisms.

Additionally, MultiMIL could be utilized for perturbation studies to understand how cells respond to various treatments or environmental changes. This application is crucial for identifying potential therapeutic targets and understanding drug response mechanisms [61]. By analyzing perturbation data, MultiMIL could reveal how different cell states shift in response to specific interventions, providing insights that can guide the development of patient-tailored drugs. This approach could potentially not only help identify effective treatments but also customize therapies to individual patients based on their unique cellular responses, thereby enhancing the precision and efficacy of medical interventions [62].

Several approaches for unsupervised sample representation learning have been developed in recent years [63, 64]. We envision MultiMIL as a complementary approach to such methods, allowing researchers to first identify subclusters of patients in an unsupervised manner and then utilize the clustering assignments to identify cell states driving the separation beyond known metadata. As an extension, we experimented with incorporating sample-level covariates such as sex and age by encoding them and providing them to the model. This did not lead to performance gains, suggesting that simply appending covariates may not be sufficient. Nevertheless, we see substantial potential in this direction; future research will need to investigate how covariates should be represented, where they should be introduced in the architecture, and what interactions with cellular features are most informative.

MultiMIL offers a novel approach for linking single-cell-level and sample-level data, identifying bi-ologically meaningful disease-associated cell states. By accommodating raw data and embeddings from existing atlases or foundation models, our method provides computational biologists with a versatile and scalable tool for multi-scale analysis.

## 4 Conclusions

MultiMIL is a deep-learning framework that bridges the gap between single-cell measurements and sample phenotypes through attention-based multiple-instance learning. Our model learns to pre-dict disease phenotype by prioritizing the cell states that drive the disease. This approach can be utilized with a variety of modalities and input data types, ranging from IMC data to embeddings from foundation models. We demonstrated how MultiMIL can be applied to COVID-19, idiopathic pulmonary fibrosis and breast cancer datasets, not only recovering known disease states but also expanding molecular signatures. MultiMIL is a scalable and interpretable tool for linking single-cell measurements and patient phenotypes, enabling multi-scale learning in single-cell genomics.

## 5 Declarations

### Ethics approval and consent to participate

Not applicable.

### Consent for publication

Not applicable.

### Availability of data and materials

The package is available at http://github.com/theislab/multimil and the pipeline for sample-level prediction is available at https://github.com/theislab/sample-prediction-pipeline. The code to reproduce the results and figures is available at http://github.com/theislab/multimil_reproducibility. All datasets analyzed in this manuscript are public and can be downloaded through http://github.com/theislab/multimil_reproducibility.

## Authors’ contributions

FJT conceived the project with contributions from ML and AL. AL, ML and FJT designed the model. AL implemented the model. AL performed the benchmarks. AL, SHZ and AAM analyzed cell attention scores for the PBMC case study. AL performed the analysis for the HLCA and the breast cancer case studies. All authors contributed to the manuscript. ML and FJT supervised the project.

## Acknowledgements

We thank Dr. Luke Zappia for the figure and manuscript feedback, Jan Engelmann and Vladimir Shitov for discussions on MIL and patient representation learning, Maiia Shulman for discussions on breast cancer, Dr. Daria Romanovskaia for the manuscript feedback and Dr. Malte D. Luecken for the constructive feedback throughout the project. We thank the scverse community (especially the developers and the maintainers of scanpy and scvi-tools packages) and the Theis lab for valuable discussions.

## 6 Methods

### MultiMIL training

MultiMIL is a deep learning model that consists of a pooling layer and a feed-forward classification network. We assume the input to the model is single-cell embeddings, where assignments of cells to samples and sample-level labels are known. We will now focus on a single mini-batch and describe one forward pass of the model. Each training batch consists of single-cell data from *l* samples {*X*_1_*, . . ., X_l_*} and the sample labels {*p*_1_*, . . ., p_l_*}. The number of rows in each matrix *X_i_* equals *n*, which is the number of cells in the sample mini-batch, and the number of columns equals *h*, which is the number of features in the input space.

The single-cell representations are first fed into the pooling layer. The goal here is to aggregate the representations of all cells *z* ∈ R*^n^*^×*h*^ from the sample (i.e., bag) into a *z*_bag_ ∈ R*^h^*. This bag representation corresponds to a pooled representation of a bag of cells. The pooling functions are applied independently to each latent dimension, so we will refer to the representation of cell *i* along latent dimension *p* as *z^p^* ∈ R and explain how the pooled representation is calculated for one dimension. There are several ways to obtain this pooled representation, e.g., applying *max* operator

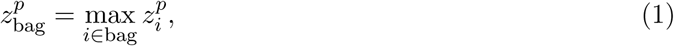

*mean* operator,

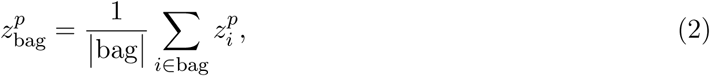

where ^1^bag^1^ indicates the number of cells in a bag. More advanced pooling functions include attention or gated attention pooling [15], where first a scalar value *a^i^* ∈ R per cell is learned with an attention mechanism [65]:

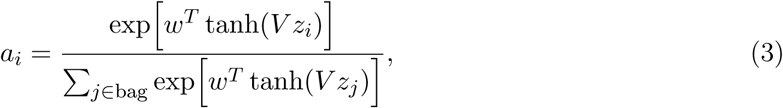

or gated attention mechanism [66]:

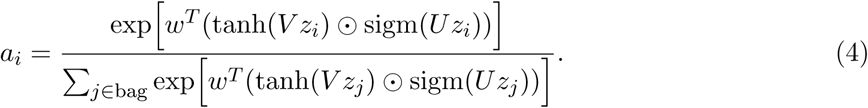

Then the bag representation is learned as a weighted average:

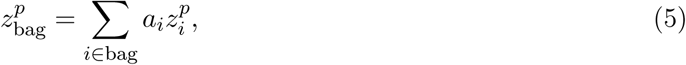

where *w* ∈ ℝ*^q^, V* ∈ ℝ*^q^*^×*h*^ and *U* ∈ ℝ*^q^*^×*h*^ are learnable weights and *q* is a hyperparameter known as attention dimension. We note that the weight *a_i_* is learned per cell and not per cell per latent dimension.

After the aggregation, *z*_bag_ is fed into a classification network that consists of blocks of a linear layer with dropout, layer normalization and nonlinearity. The number of neurons in the last layer equals the number of classes for the classification tasks. The classification network predicts the distribution of disease labels for a given bag (i.e., sample). For regression tasks, the last layer consists of one neuron and predicts a scalar value per bag. Next, we will discuss the training loss.

The classification loss is calculated as the cross-entropy loss between one-hot encoded true sample labels and the predicted values of the final layer in the classification network. If the user is interested in modeling the sample labels as a progression, the last layer of the classification network can be changed to a regression head. In this case, the regression loss is calculated as mean squared error loss. To calculate the accuracy of the per-sample predictions, we round the prediction to the closest true value. For simplicity, we refer to the regression loss also as the classification loss.

### MultiMIL inference

During test time, we aim to predict the disease class for new patients. We assume that the single-cell representations have the same dimensionality as the train data and that they were obtained from either mapping the query data onto the same atlas/reference with scArches [67], from the same foundation model or were left out from the train dataset as the test data. Then, we perform one forward pass through the model as described in the previous section. The pooling layer aggregates the cell representations into a bag representation, which is then classified using the classification network.

### Integration metrics

To assess the quality of the integration, we used several metrics from the scIB package [23]. Note that scIB metrics were designed for unimodal integration, and not all of them can be easily applied in the multimodal case; hence, we chose the metrics that only require the integrated embedding space as input (and not, e.g., the original unintegrated space). In the following, we briefly discuss two metrics for batch removal and four for biological variance conservation. As in scIB, the final score was calculated as 0.4*batch correction + 0.6*biological conservation. For more details on the metrics and the implementation, see [23].

### Batch correction

Graph connectivity measures how well cells from each cell type are connected in a k-nearest neighbor graph. If the connectivity is high, then the batch effect was removed sufficiently. Average silhouette width (ASW) compares average distances within a cluster with distances to other clusters. The resulting score reflects how compact the clustering is. For ASW batch, we expect the batch clusters to be well-mixed together for a high batch correction score.

### Biological variance conservation

Adjusted Rand Index (ARI) and Normalized Mutual Information (NMI) evaluate how well the clustering is aligned with the ground truth labels, i.e., cell type annotations. ASW label is a modification of ASW batch, where we expect the cell type clusters to be compact and separate from other cell type clusters for a high biological conservation score. Isolated label ASW assesses how well rare cell types are distinguishable from the rest of the data.

### Benchmarks

#### Obtaining the cell embeddings

For the hyperparameter benchmark, we first obtained the scVI, PCA, GeneFormer and scGPT embeddings of the full Stephenson et al. dataset [20]. scVI was trained with the default parameters with ’Site’ and ’sample_id’ as batch covariates. We used helical [68] to obtain embeddings from GeneFormer (*gf-12L-95M-i4096* model) and scGPT, both in a zero-shot setting. We note that scVI embedding has 30 latent dimensions, PCA has 50, and GeneFormer and scGPT both have 512.

#### Benchmakring hyperparameters

We reported the accuracy and the standard deviation of MultiMIL classification and regression models on the top five performing models for each of the embeddings. The tested hyperparameters are shown in Table 1.

To assess the effects of learning rate, batch size and the pooling function, we focused on the scVI and scGPT embeddings as they scored best with respect to scIB scores. Hence, we reported across all runs for these two embeddings.

#### Classification prediction

We compared MultiMIL to several baselines: random forest, multiclass logistic regression and feed-forward neural networks. We trained each model on the following data input types: mean embeddings, cell-type mean embeddings, cell-type frequency vectors and cell embeddings. We note that some baselines, namely cell-type mean embeddings and cell-type frequency vectors, require cell-type information, while MultiMIL and the rest of the baselines are entirely unsupervised.

The benchmark was performed on the Stephenson et al. PBMC dataset [20]. Throughout these experiments, we used the scVI embeddings. We created 5-fold cross-validation splits based on patient information, i.e., so that cells in each train/validation split come from different patients. We used sklearn.model_selection.KFold() to create the splits and sklearn.metrics.classification_report() to report the classification accuracy. We additionally optimized the learning rate for the IMC dataset [22] as the feature set and the measurement distribution are substantially different from those of single-cell data.

The parameters tested in the grid search are shown in Table 1.

Depending on the batch size, the number of samples in each batch varied from 2 to 8, as 128 cells from each sample were drawn for the forward pass. The default parameters were chosen based on the prediction accuracy of the validation set averaged across five splits.

Next, we discuss the baseline models and the input data in more detail. We performed a hyperparameter grid search for NN-based models and reported the best-performing configuration. Patient disease labels were used as class labels throughout this benchmark apart from the “Cell embedding” input type, where all the cells from a diseased donor were assumed to have the disease class.

#### Baseline models

- Multiclass logistic regression is an extension of the logistic regression method that allows the prediction of multiple classes. We calculate the probability of belonging to a particular class with a softmax function and calculate the loss as the entropy between predicted probabilities and the true class. We optimize the loss function with gradient descent.
- Random forest was implemented using sklearn.ensemble.RandomForestClassifier() with the default parameters.
- Neural network was implemented as a 2-layer feed-forward network with one hidden layer of 64 neurons, batch normalization and ReLU activation. The second linear layer outputs class probabilities. We trained the neural network baselines with Adam optimizer [69] for 200 epochs for sample-level inputs and 30 epochs for cell-level inputs. Hyperparameter search was run for batch size and learning rate shown in Table 2.

#### Input data types

- Mean embedding representations were calculated from the latent embeddings with decoupler.get_pseudobulk() specifying the sample parameter and keeping all the cells.
- Cell type-aware mean embedding representations were calculated from the latent embeddings with decoupler.get_pseudobulk() specifying the sample and group (i.e., cell type) parameters and keeping all the cells. To obtain one representation per sample, we concatenated cell-type-specific vectors into one vector.
- Frequency vectors were calculated from cell type proportions for each sample.
- Low-dimensional cell embeddings were directly passed to the baselines, e.g., from atlases or foundation models.

#### Identification of DA cell states

We ran the default Milo [13] analysis on the PBMC dataset using the embeddings from scVI following the pertpy [70] implementation. We ran three pairwise analyses comparing mild COVID-19 and healthy, severe COVID-19 and healthy, and severe and mild COVID-19. We show the neighborhoods with spatial false discovery rate (FDR) corrected levels of less than 0.05.

**Table 2.** Hyperparameter search NN baseline.

| Hyperparameter | Description | Range |
| --- | --- | --- |
| Learning rate | learning rate parameter | $\{1e-5, 1e-4, 1e-3\}$ |
| Batch size for sample-level inputs | size of the training mini-batch | $\{8, 16, 32, 64\}$ |
| Batch size for cell-level input | size of the training mini-batch | $\{128, 256, 512, 1024\}$ |

**Table 3.** Running times for MultiMIL.

|  | average runtime (s) | standard deviation (s) |
| --- | --- | --- |
| classification setting | 356 | 9 |
| regression setting | 357 | 5 |

**Table 4.** Default architecture for MultiMIL.

|  | Layer |
| --- | --- |
| MIL attention pooling | calculation of attention scores as in Eq. 4<br>calculation of the weighted sum as in Eq. 5 |
| Classifier network | Linear(embed_dim, 128)<br>Dropout(0.2)<br>LayerNorm<br>LeakyReLU<br>Linear(128, n_classes) |

To identify cell types with the highest compositional change between healthy and severe (**Fig. 3e**), we excluded the cell types with fewer than 1,000 cells and calculated the relative change in proportions between the number of cells in pooled severe and healthy samples.

#### Sample representations

We also investigated how well we can predict sample labels with a kNN classifier from sample representations obtained by aggregating cell embeddings with different aggregation strategies. We used the attention scores from the best-performing model trained on the scVI embedding throughout these comparisons. Sample representations were calculated as either the mean of cell embeddings belonging to the sample or the weighted average (where the weight for each cell was the learned attention score). We used cells with top 10%, 20%, 30%, 40%, 50%, 60%, 70%, 80%, 90% highest attention score for the aggregation.

#### Calculation of the profibrotic signature

To calculate the profibrotic score for macrophages in HLCA, we used the signature from [29]: *SPP1, LIPA, LPL, FDX1, SPARC, MATK, GPC4, PALLD, MMP7, MMP9, CHIT1, CSTK, CHI3L1, CSF1, FCMR, TIMP3, COL22A1, SIGLEC15, CCL2*. The score was calculated with scanpy.tl.score_genes(). We performed a two-sided t-test to check for the significance of the score in all IPF macrophages vs. IPF macrophages with the high attention score using scipy.stats.ttest_ind(). We used edgeR-QLF [12] to identify the genes differentially expressed in IPF macrophages with high attention compared to all IPF macrophages and reported genes with a log-fold change greater than 1.5 and FDR-corrected p-value less than 0.01 as up-regulated (see Supplementary Table 1).

#### Gene Ontology analysis

We used GOATOOLS [71] to run the GO term analysis on the genes that were identified as signif-icantly upregulated in the IPF macrophages with MultiMIL. We followed the tutorial and ran all the functions with their default parameters. We reported the terms with the corrected p-value less than 0.1 as significant.

#### Identification of differential signal in IMC data

We performed a Wilcoxon rank-sum test between the high-attention (top 10%) tumor cells, compar-ing cells from each of the three grades with the rest of the high-attention tumor cells, and reported the Benjamini-Hochberg corrected p-values.

#### Datasets

Stephenson et al.

The PBMC dataset contains 647,366 cells from 130 donors, collected at three sites. All scRNA-seq data points were used for the integration. For the prediction experiment with all COVID-19 stages, we removed non-COVID and non-healthy samples, so the prediction was performed for 6 classes: healthy (23 samples), asymptomatic (9), mild (23), moderate (30), severe (13) and critical (15). For the 3-class prediction, we subset the data to 13 healthy, 13 mild and 13 severe samples to perform the benchmarking experiments on the balanced data and not confound the performance with unbalanced classes. For the binary experiment, the COVID-19 class had 90 samples and the healthy class had 23 samples. We used ’sex’ and ’age interval’ as categorical sample covariates in the experiments where we concatenated one-hot covariates to the sample embeddings before passing them through the classification network.

Sikkema et al.

Human Lung Cell Atlas (HLCA) consists of the core (584,444 cells, 107 donors) and the extension datasets (1,797,714 cells, 380 donors). The core samples are all healthy, while the extension has healthy and diseased samples. In our experiments, we subset the data to healthy and IPF samples in a balanced way, i.e., the number of donors is the same (67) in both groups.

Jackson et al.

The human breast cancer dataset consists of two cohorts, the Zurich and the Basel cohorts. In our experiments, we subsetted the data to the Zurich cohort and excluded metastatic samples from the analysis. This dataset consists of 72 donors, 347 samples and 395,769 cells. We used ’core’ as the sample covariate in our experiments. We trained the model to predict grades 1 (95), 2 (122) and 3 (130).

#### Running time

We provide training times for MultiMIL with the default architecture in Table 3. The training was performed on the same GPU server with the following characteristics: Intel(R) Xeon(R) Platinum 8280L CPU with 28 cores, 2.70GHz, Tesla V100-SXM3-32GB GPU. We report the average run time and standard deviation across three runs. We used the PBMC CITE-seq dataset [20], subsetted to healthy, mild and severe COVID-19 in a balanced way, resulting in 256,051 cells. All models were trained for 50 epochs. We modeled the prediction task as either a three-class classification problem or a regression problem.

#### Default architecture

The default architecture for the classification setting of MultiMIL is shown below. In the regression setting, the last layer has only one output neuron.

We note that embed_dim refers to the number of latent dimensions in the low-dimensional representation of single-cell data from atlases or foundation models, or the raw dimensionality of the input if raw data is used. For the model versions, where we incorporated one-hot encoded sample covariates, the embed_dim equals the number of latent dimensions in the low-dimensional representation of single-cell data plus the number of categories for each of the covariates. For instance, if we concatenate the sex covariate with two categories present, one-hot encoded embeddings will add two dimensions to embed_dim.

---

**Supplementary Figure 1.**
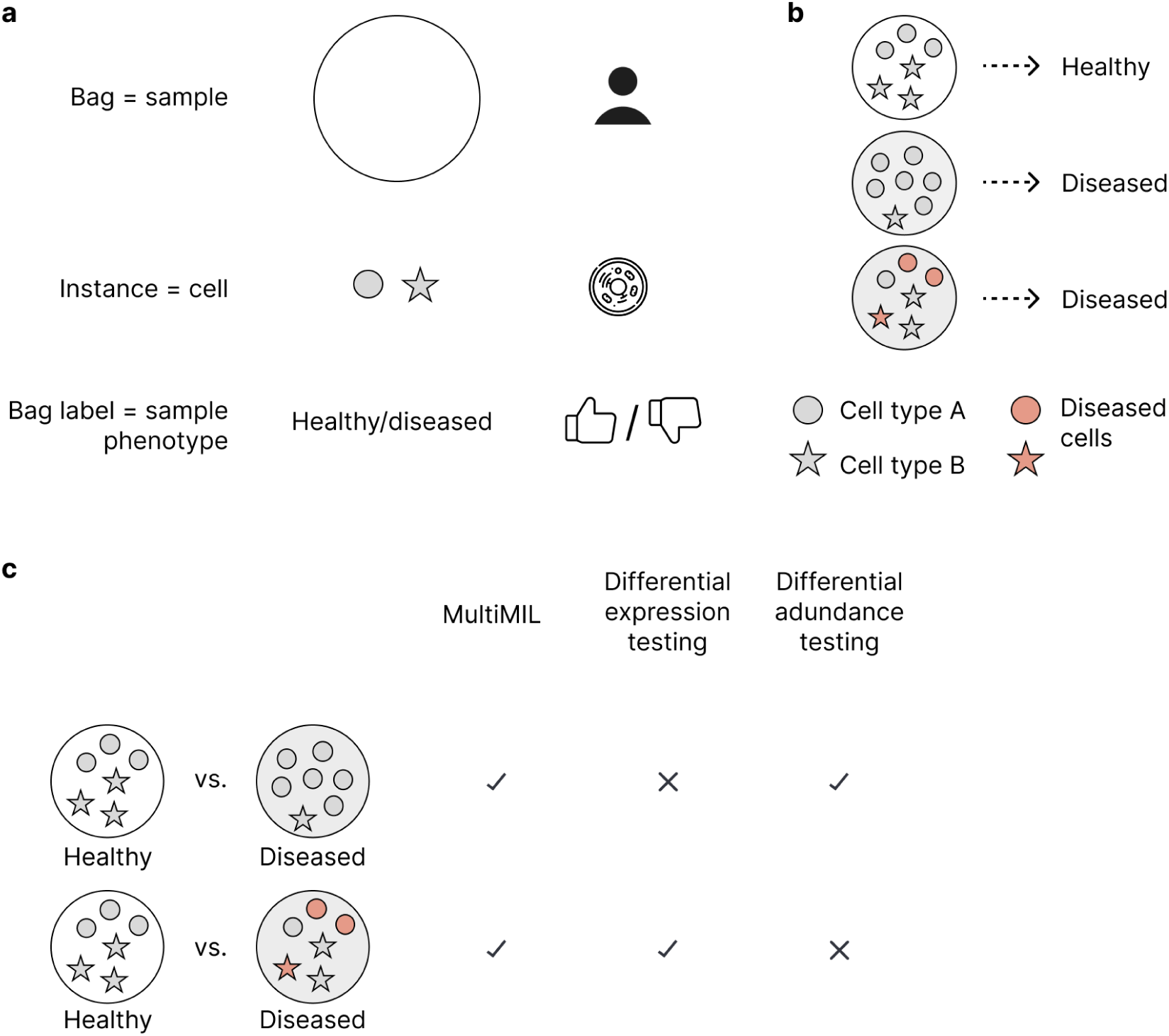
Multiple instance learning. **(a)** In our context, bags correspond to samples, instances to cells and the classification labels are known for bags. **(b)** Examples of data points in the multiple-instance-learning dataset. Our task is to classify bags into classes and identify cells (i.e. colored instances) that are associated with a certain disease. **(c)** MultiMIL can identify changes in the abundance of cell types between conditions as well as transcriptomic changes.

**Supplementary Figure 2.**
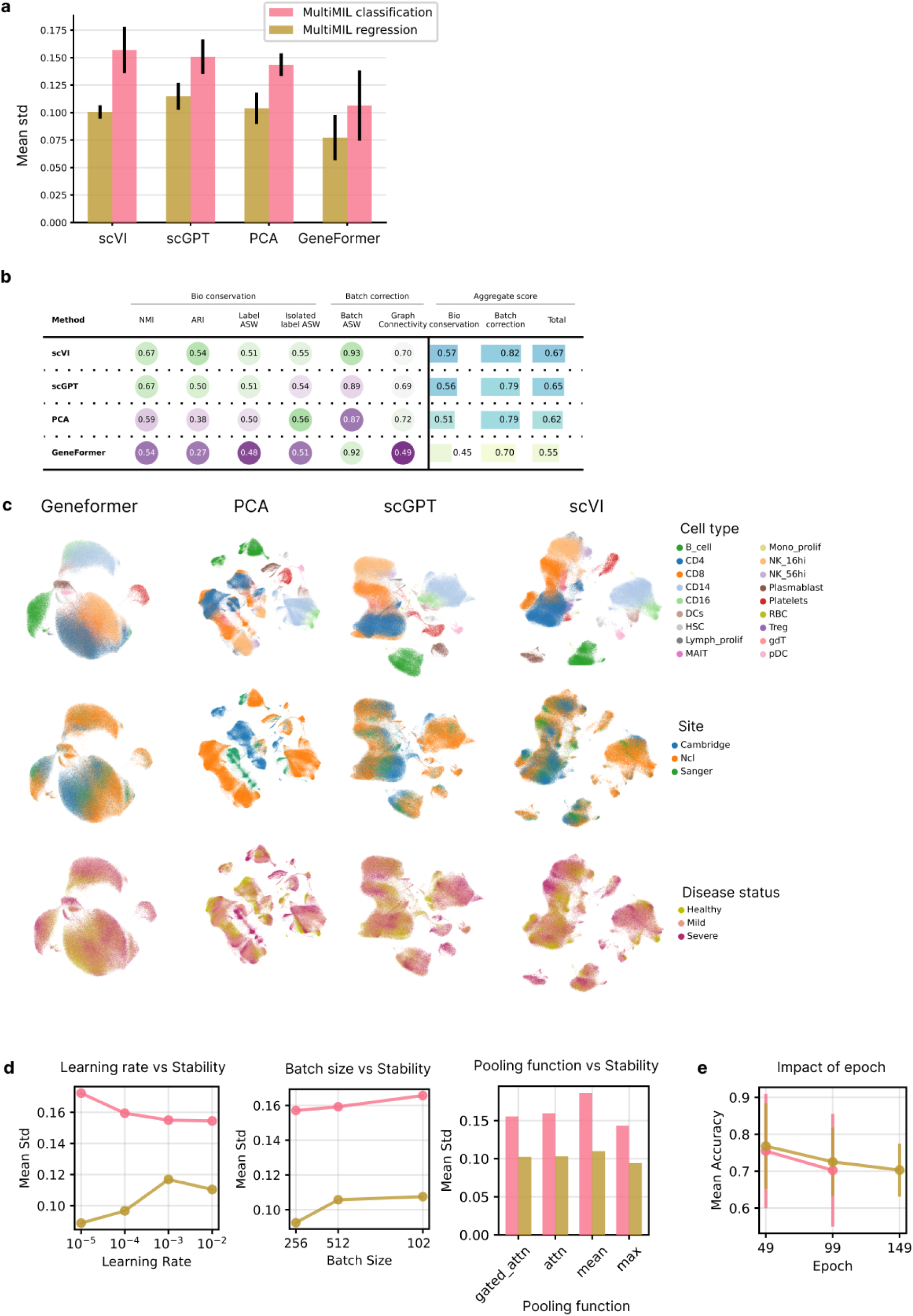
Benchmarking MultiMIL on different embeddings for Stephenson et al. **(a)** A bar plot showing the mean standard deviation of prediction accuracy across cross-validation splits for scVI, PCA, scGPT and GeneFormer embeddings, taking the top 5 MultiMIL models, colored by regression/classification setting. **(b)** scIB metrics assessing the quality of the embeddings. **(c)** UMAPs of the GeneFormer, PCA, scGPT and scVI embeddings, colored by cell type, batch (i.e., site) and disease stage. **(d)** Plots showing the mean standard deviation of prediction accuracy depending on the learning rate, batch size and the pooling method, colored by MultiMIL’s regression/classification setting. **(e)** Effect of the epoch, i.e., how long the model was trained for, on the accuracy.

**Supplementary Figure 3.**
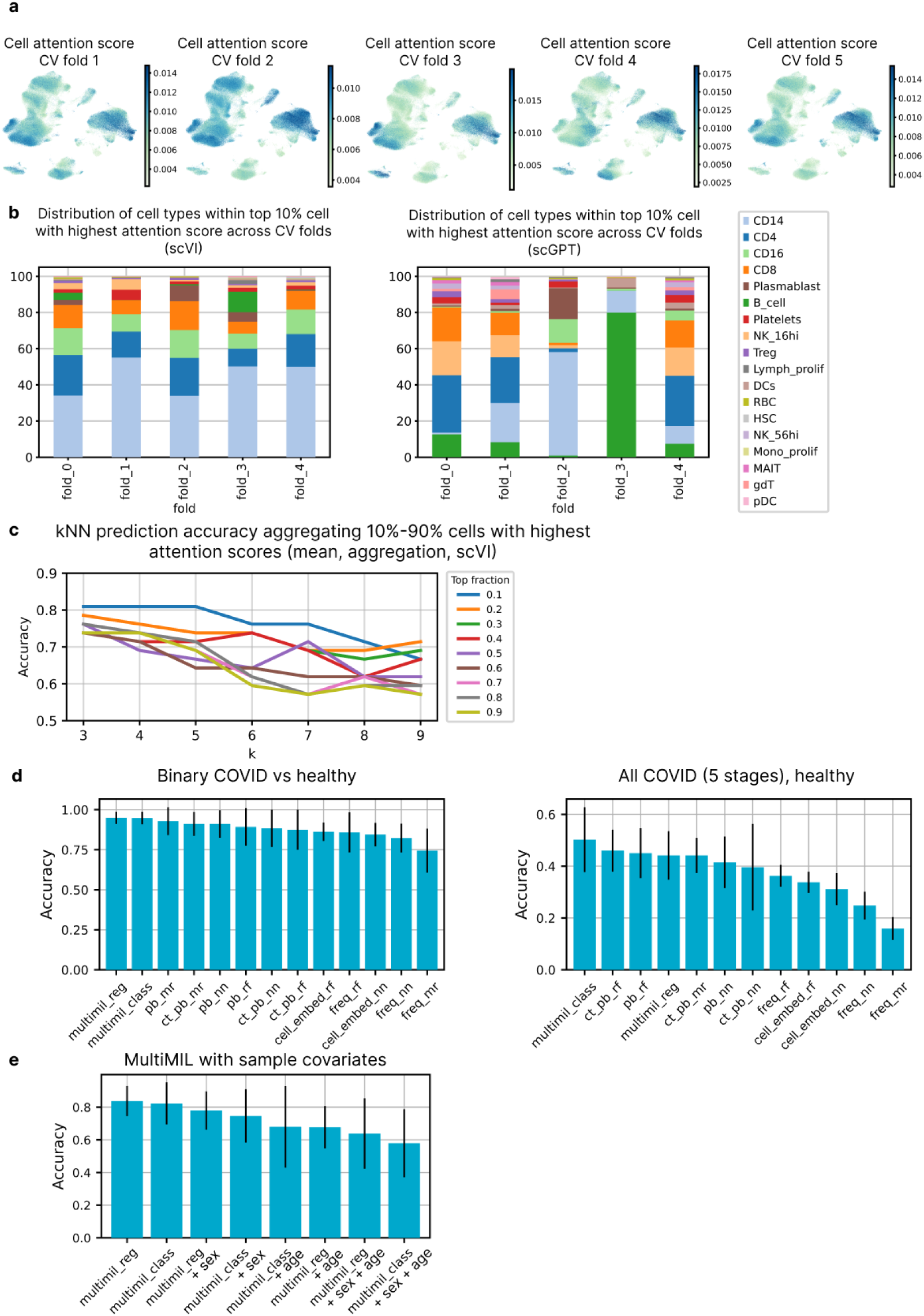
MultiMIL’s cell attention scores, sample representations and prediction accuracy across different tasks. **(a)** UMAPs showing cell attention scores learned in five cross-validation runs using scVI embedding and MultiMIL’s regression setting. **(b)** Stacked bar plots showing the distribution of cell types with top 10% highest attention scores across five cross-validation runs, using top-performing MultiMIL’s regression models on scVI and scGPT embeddings. **(c)** Line plots showing how well the kNN classifier can predict sample labels from 3, 4, 5, 6, 7, 8, 9 nearest neighbors when the sample representation was obtained by averaging cell embeddings of cells with top 10%-90% highest attention scores. **(d)** Results of the prediction benchmark on binary (healthy, COVID-19) and full data (healthy, 5 COVID-19 stages) using scVI embeddings, comparing MultiMIL with the baselines. **(e)** Results of the prediction benchmark on the three-class (healthy, mild and severe COVID-19) task using scVI embeddings, comparing MultiMIL default models with MultiMIL models incorporating sample covariates (sex and age).

**Supplementary Figure 4.**
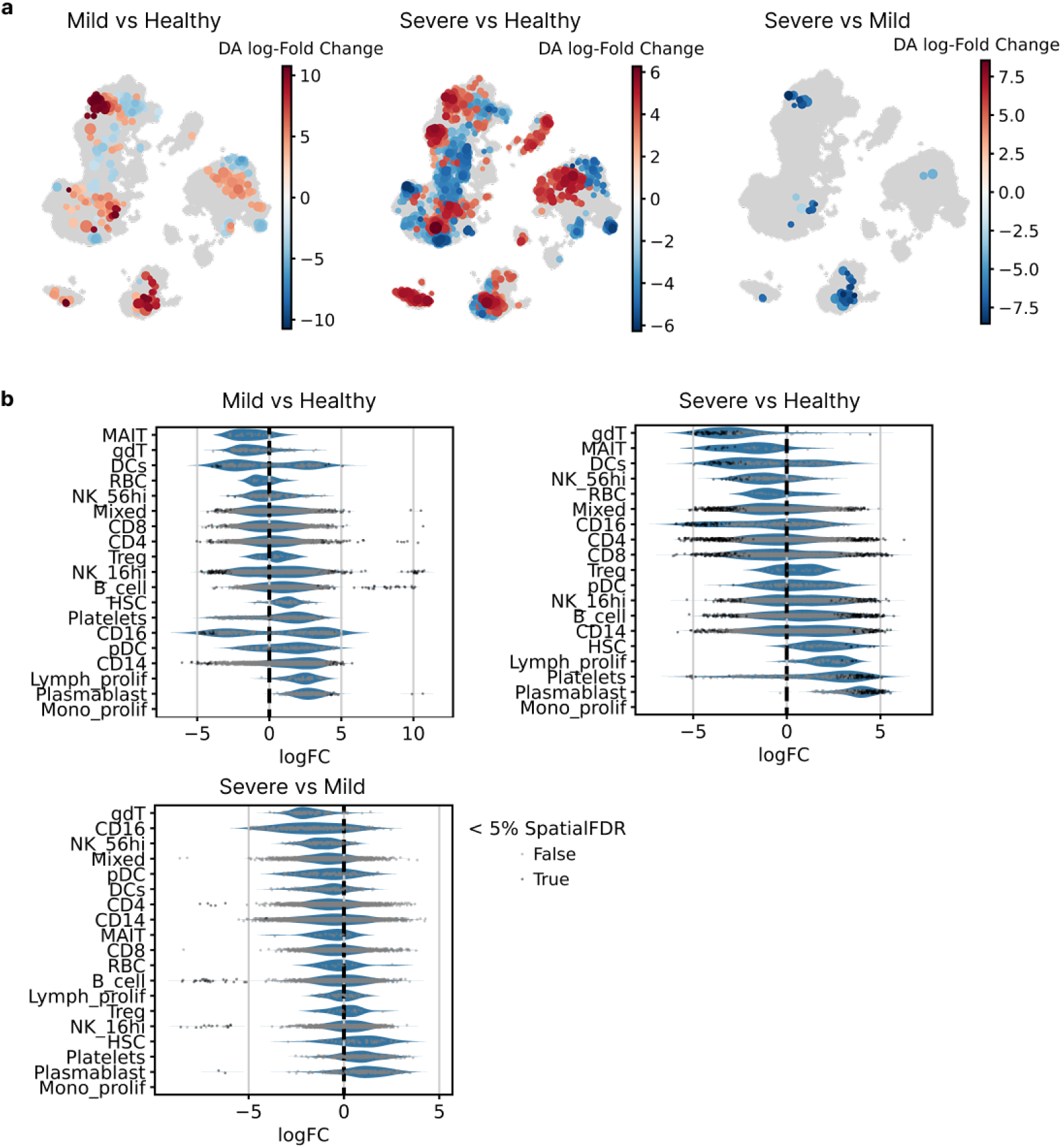
MILO analysis of Stephenson et al. dataset. **(a)** Results of Milo analysis run on MultiMIL’s embeddings, mild vs. healthy, severe vs. healthy and severe vs. mild, each colored by DA log-fold change (red corresponds to the first condition in the title). **(b)** Violin plots showing DA changes for each of the cell types in mild vs. healthy, severe vs. healthy and severe vs. mild.

**Supplementary Figure 5.**
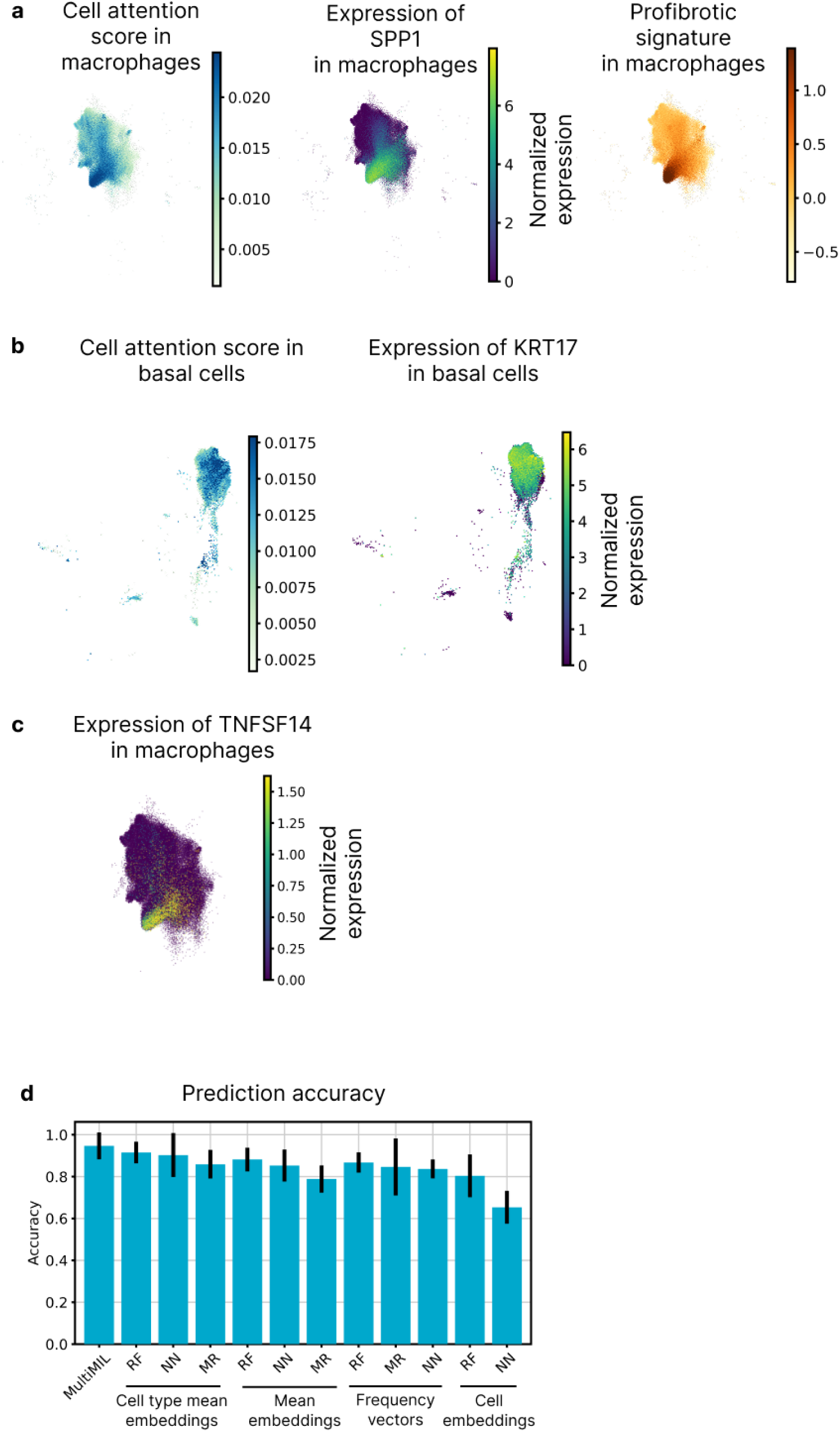
IPF in HLCA. **(a)** UMAPs of macrophages, colored by cell attention, normalized expression of SSP1 and profibrotic signature score. **(b)** UMAPs of basal cells, colored by cell attention and normalized expression of KRT17. **(c)** The UMAP of macrophages, colored by the normalized expression of TNFSF14. **(d)** Prediction accuracy of MultiMIL compared to other baseline models on the binary classification task, i.e., IPF vs healthy.

**Supplementary Figure 6.**
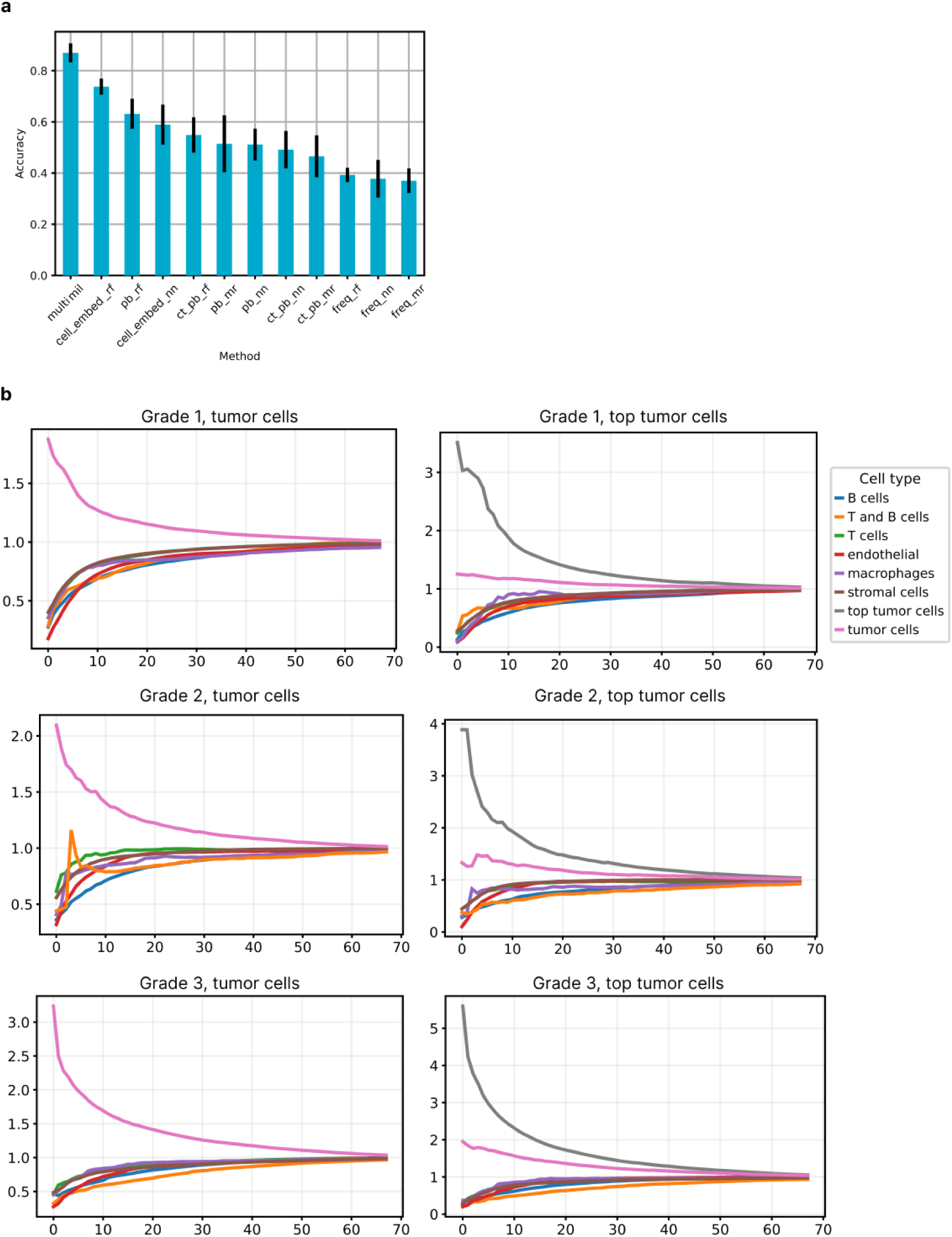
Predicting grades in breast cancer from IMC data. **(a)** Prediction accuracy of MultiMIL compared with baselines. **(b)** Co-occurrence analysis of tumor cells (left columns) and high-attention tumor cells (i.e., top tumor cells), split by grade.

